# Haplotype and diversity signatures of ultra-soft selective sweeps in HIV-1

**DOI:** 10.64898/2026.07.21.739925

**Authors:** Elena V. Romero, Dylan A. Clark, Lillian B. Cohn, Alison F. Feder

## Abstract

Many urgent medical and agricultural challenges are driven by resistance evolution via soft selective sweeps of multiple simultaneous mutations. Standard approaches to detect these mutations involve genome scans for regions with reduced diversity and increased haplotype lengths. However, it is unknown the extent to which those signatures persist as the number of mutations driving resistance grows. Here, we analyzed longitudinal linkage-resolved data from 10 intra-host HIV populations treated with broadly neutralizing antibody 10-1074. We found that HIV escapes 10-1074 with minimal perturbations to diversity and haplotype homozygosity in the region surrounding the sweep in the majority (8/10) of treated individuals. We matched these *in vivo* escape trajectories to forward simulations and found that adaptive mutations conferring escape must have been present on 20 or more genetic backgrounds to generate these signatures. These “ultra-soft” sweep signatures more closely resemble genetic patterns in a treatment non-responder without an adaptive response to 10-1074 than those of two other trial participants where adaptation occurred via harder selective sweeps. Our results demonstrate that HIV can adapt to a broadly neutralizing antibody treatment while retaining nearly all of its standing genetic diversity and that selection scans dependent on regional diversity and haplotype homozygosity signatures fail in this “ultra-soft” regime.

## Introduction

In their seminal works on the topic, Pennings and Hermisson demonstrated that when populations produce beneficial mutations at a sufficiently high rate, multiple of these beneficial mutations should spread simultaneously as “soft” selective sweeps [Pennings and Hermisson, 2006a, Hermisson and Pennings, 2005, 2017, Pennings and Hermisson, 2006b]. Adaptation via soft selective sweeps underlies some of the world’s most urgent medical and agricultural challenges, including antimicrobial, pesticide and herbicide resistance [Nair et al., 2007, Délye et al., 2013, Pennings et al., 2014, Romero et al., 2026, Messer and Petrov, 2013]. Because beneficial mutations land on multiple genetic backgrounds that are selected to high frequency, soft selective sweeps leave more modest genetic signatures than classical hard sweeps, including an attenuated reduction of genetic diversity (Figure 1). As a result, new methods to specifically identify soft sweeps signatures in genomic data often focus on more robust patterns related to haplotype structure rather than diversity [Garud et al., 2015, Schrider and Kern, 2016, Harris and DeGiorgio, 2020, Zhao et al., 2024]. These methods have revealed that soft sweeps are more widespread than was previously understood.

**Figure 1:**
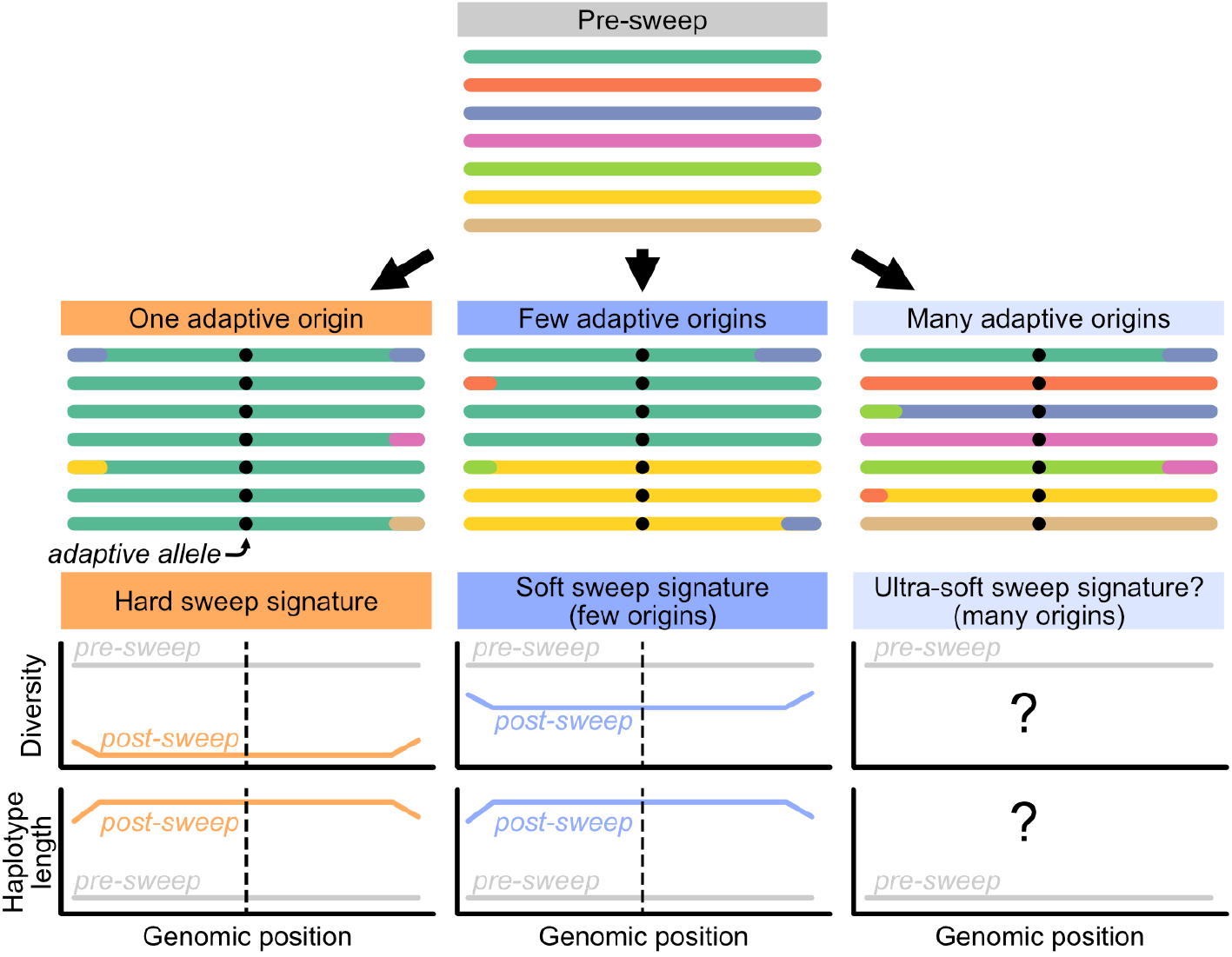
Signatures for sweeps of one, few or many adaptive origins. Line diagrams represent genomes colored by haplotype at pre-sweep and post sweep sampling times for three different sweep scenarios. Cartoons of diversity and haplotype length show how these metrics compare across sweep scenarios.

Despite the growing appreciation for the prevalence of soft selective sweeps, most empirical, theoretical and methodological work on the topic has focused on cases where the number of origins of adaptive mutations is small (i.e., *<* 10) rather than large (although see Kreiner et al. [2019], Anderson et al. [2017], Cao et al. [2025]). Multiple factors contribute. First, estimates of effective population sizes can suggest that beneficial mutation input (i.e., *N_e_× µ_b_* or the number of beneficial mutations entering the population each generation) may be too limited to generate more than a few origins of adaptive alleles. However, phenotypes like resistance often emerge in populations with extremely large census population sizes (pathogens, insects, plants), where long-term estimates of *N_e_*may not accurately reflect how many adaptive mutations enter a population each generation in the short-term [Messer and Petrov, 2013]. For example, estimates for HIV intra-host effective population size using long term coalescent expectations suggest that *N_e_ ≈* 1000 [Brown, 1997], while estimates based on the rate of production of adaptive alleles are two orders of magnitude higher [Pennings et al., 2014]. Second, most clear instances of soft sweeps described in the literature have few origins of resistance [Garud et al., 2015, Tishkoff et al., 2007, Pennings et al., 2014]. These empirical observations have concentrated theory and methods on understanding and identifying these cases. However, soft sweeps of few origins may simply be easier to identify because they generate characteristic extended haplotype signatures, while sweeps with more origins have not been well-characterized and may leave subtler patterns. Together, these points suggest that soft sweeps of many beneficial mutations could drive adaptive evolution, but we may fail to recognize them because of their unknown genetic signatures.

We recently analyzed *in vivo* HIV intra-host evolution during escape from broadly neutralizing antibody 10-1074 monotherapy and found multiple lines of evidence of soft selective sweeps from a large number of origins [Romero et al., 2026]. Viral populations in each of 10 participants acquired 4–13 unique amino acid changes in the HIV Env protein which are each sufficient to confer escape [Radford and Bloom, 2025]. These mutations were often present on multiple haplotypes, and were occasionally encoded by multiple nucleotide mutations that generated the same amino acid change. Deep, haplotype-resolved resequencing data of the HIV *env* gene provided an opportunity to understand the genetic diversity and haplotype signatures associated with these selective sweeps of many origins and how they differ from sweeps with fewer origins.

In this paper, we found evidence that the sweeps of intra-host HIV adaptation to broadly neutralizing antibody 10-1074 minimally impact post-sweep diversity adjacent to the site under selection and can increase diversity at the selected site itself. This strongly contrasts expectations under classical sweeps. More strikingly, most (8/10) intra-host populations also did not exhibit extended haplotype homozygosity surrounding the selected loci, despite haplotype signatures being the basis of most tests for soft selective sweeps. We call this pattern an “ultra-soft” selective sweep. To contextualize these unusual findings, we simulated sweeps with large numbers of artificially introduced highly beneficial mutations. We found that these simulations replicate the *in vivo* diversity and haplotype homozygosity we observe during 10-1074 escape when the number of introduced beneficial mutations is sufficiently large. Comparing between the *in vivo* data and our simulations, we estimated that HIV escape from 10-1074 is likely driven by beneficial mutations on *≥* 20 backgrounds. Collectively, these analyses reveal that some selective sweeps may be so soft as to be effectively invisible in genetic data.

## Results

### HIV escapes broadly neutralizing antibody 10-1074 via ultra-soft sweeps in human trial data

Caskey et al. 2017 profiled viral escape in 33 people living with HIV following the administration of 10-1074, a broadly neutralizing antibody targeting a conserved glycan in the V3 loop region of the HIV envelope glycoprotein (Env) [Caskey et al., 2017]. Recently, Romero et al. [2026] performed deep resequencing from ten of these individuals who experienced initial viral load suppression followed by rebounds in viral load within 10 weeks (Figure 2A). This sequencing revealed that viral escape from 10-1074 across this cohort was driven by soft selective sweeps of multiple amino acid substitutions at Env positions 325, 332 and 334 which prevent 10-1074 binding (Figure 2B). These escape mutations were identified because they disrupted known 10-1074 contact sites [Mouquet et al., 2012, Garces et al., 2014, Gristick et al., 2016] and were confirmed via *in vitro* antibody neutralization assays [Caskey et al., 2017]. Mutations at sites 332 and 334 cause the loss of the glycosylation site (N332) while mutations at site 325 disrupt a nearby 10-1074 contact site [Bricault et al., 2019, Caskey et al., 2017, Dingens et al., 2019, Radford and Bloom, 2025]. These mutations were at low or undetectable frequencies pre-treatment and only rose following the application of therapy [Romero et al., 2026]. While both studies discovered clear evidence of adaptation, and that multiple mutations drove escape from 10-1074 simultaneously, neither study examined the associated diversity and haplotype homozygosity signatures following escape. Here, we use these two datasets to quantify the signatures of extremely soft selective sweeps.

**Figure 2:**
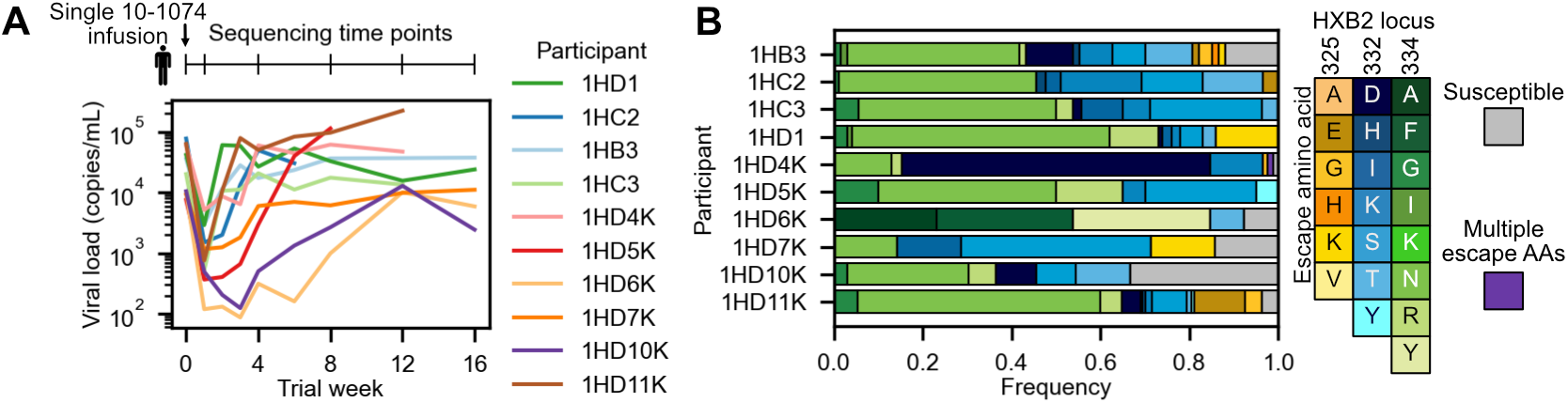
Intra-host HIV escapes 10-1074 via multiple distinct amino acid changes. **(A)** (Top) Trial schematic and sequencing time points for 10-1074 *in vivo* trial (NCT 02511990) [Caskey et al., 2017]. (Bottom) Viral load trajectories throughout the 10-1074 trial. **(B)** Frequencies of escape mutations at the first sampled time point after each participant’s viral load nadir (week 8 in participants 1HD6K & 1HD10K and week 4 in all others). Coloring indicates mutations at loci 325, 332 or 334, which are associated with 10-1074 escape [Bricault et al., 2019, Caskey et al., 2017, Dingens et al., 2019, Radford and Bloom, 2025]. Bars are colored based on the frequency of sequences carrying a single mutation at these loci (yellows, blues, greens), carrying multiple mutations (purple), or carrying no escape mutations at these loci (“susceptible”, gray). Note, only 1 sequence in one participant (1HD4K) carried multiple escape mutations.

### *In vivo* HIV escape from 10-1074 generates minimal diversity change at linked sites, instead increasing diversity at loci under selection

A characteristic signature of selective sweeps is a decrease in diversity in the region linked to the site under selection [Maynard and Haigh, 2007, DeGiorgio et al., 2016]. However, the magnitude of the decrease depends on the softness of the sweep (Figure 1) [Pennings and Hermisson, 2006a, Feder et al., 2016, Pennings et al., 2014]; as beneficial mutations land on more genetic backgrounds, more linked genetic diversity is preserved through the sweep. We investigated diversity throughout the 10-1074 trial to assess whether large numbers of beneficial mutations mitigated the diversity loss typically experienced during sweeps caused by strong selective forces like treatment initiation. We compared average pairwise nucleotide diversity (*π*) between sequences sampled before the administration of 10-1074 and those sampled at the closest time point post-viral load nadir (i.e., the lowest measured viral load), after 10-1074 escape mutations reached high frequencies (Figures 3, S1). Across the cohort, we observed minimal decreases in diversity, contrasting what is typically associated with selective sweeps; the average change in diversity across the full sequenced *≈* 2.5Kb region was *−*3.6% (IQR: *−*9.07% - 7.21%) (Figure S1). Intermediate frequency alleles play an outsize role in determining the average pairwise diversity within a population [Ferretti et al., 2018], and here we observed that nearly all mutations lost after treatment initiation were rare pre-treatment, explaining the minimal changes in diversity (Figure S2).

**Figure 3:**
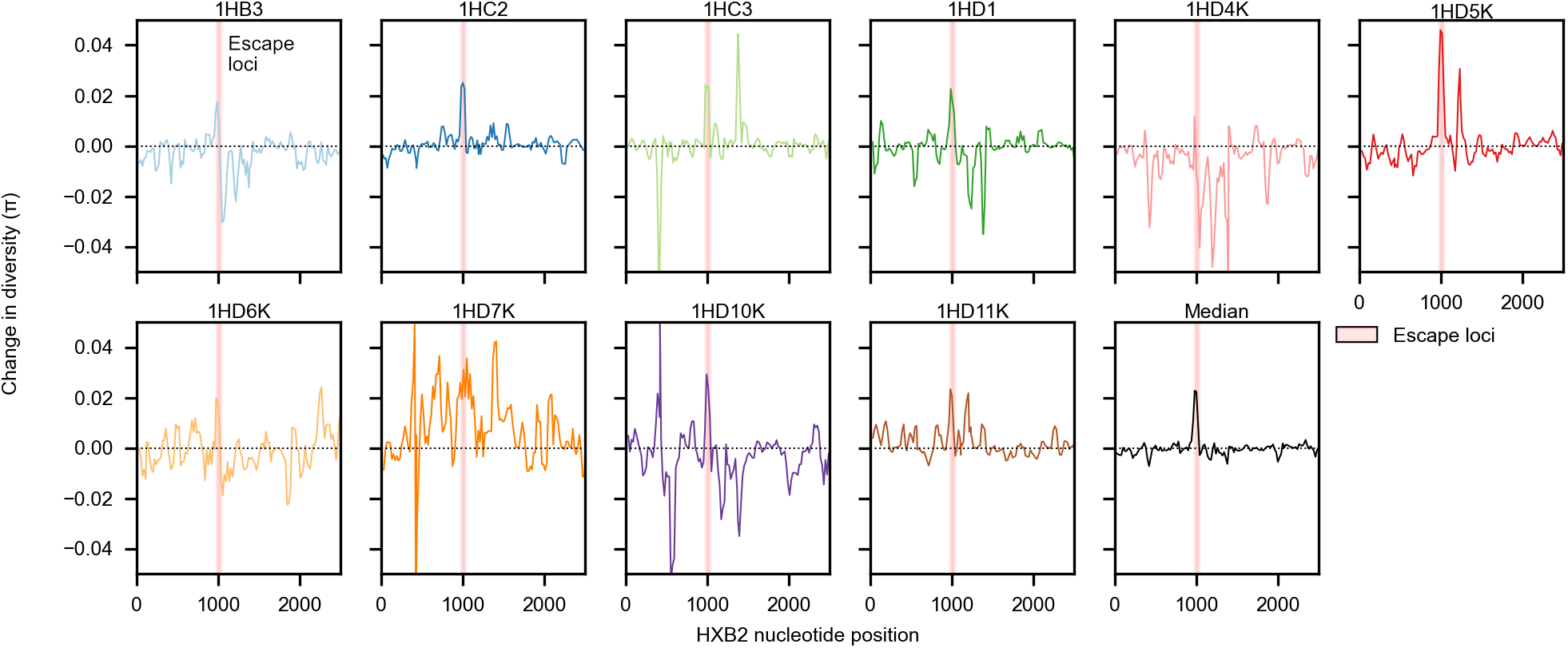
Diversity increases at escape loci following 10-1074 escape but is minimally affected in the surrounding region. Change in average pairwise Hamming distance between sequences collected prior to the 10-1074 infusion and at the first sampled time point after each participant’s viral load nadir (week 8 for participants 1HD6K and 1HD10K, week 4 for all others). Diversity change is calculated per bin based on genomic position (60 nucleotide positions per bin with bins advanced by 20 nucleotides). Each panel shows the results for an individual 10-1074 trial participant with the final panel showing the median change across all participants.

We also found little diversity loss localized in the genomic region directly adjacent to the sites under selection (Figure 3). We observed no diversity loss in the relatively conserved left flanking region of the escape locus. In the right flanking region, only participant 1HD4K showed consistent diversity loss, while participants 1HB3, 1HD1, and 1HD10K showed diversity loss which was primarily constrained the variable loops. Generally, we expect excess lost alleles in regions adjacent to sweeping alleles, since recombination has a greater opportunity to rescue genetic variation further away from the sweeping locus. However, here we observe no relationship between the probability of allelic loss and genomic distance from the selected locus (Figure S2). In further contrast to expected sweep patterns, at the selective locus itself, we observed an increase in genetic diversity in all 10 participants. This pattern is observed even in participants with decreased diversity in the flanking loci and is caused by the genetically distinct escape alleles themselves expanding after therapy initiation and differing between sequences. Collectively, HIV populations escaping from 10-1074 fail to exhibit and, in some cases, invert traditional diversity patterns: they show limited change in diversity in the selected region and increased diversity at the selected locus itself.

### *In vivo* HIV escape from 10-1074 generates little change in post-sweep haplotype lengths

A second characteristic selective sweep signature is an increase in genetic linkage [Maynard and Haigh, 2007]. As haplotypes carrying beneficial mutations rise quickly in frequency, limited opportunity for recombination generates longer linkage blocks than are typically observed under neutral evolution. Because of the relatively muted diversity signatures of soft selective sweeps and the sensitivity of genetic diversity to multiple population genetic forces, signatures based on haplotype length have proven a more robust approach to detect sweeps [Pennings and Hermisson, 2006a]. We therefore hypothesized that HIV escape from 10-1074 may still have associated haplotype signatures despite its minimal effect on diversity.

To investigate these haplotype based signatures, we calculated the integrated haplotype homozygosity (iHH) [Sabeti et al., 2002, Voight et al., 2006], a widely-used measure of haplotype length surrounding each locus. We compared the relative haplotype lengths before and after administration of 10-1074 at all variable sites in the Env gene (Figure 4).

**Figure 4:**
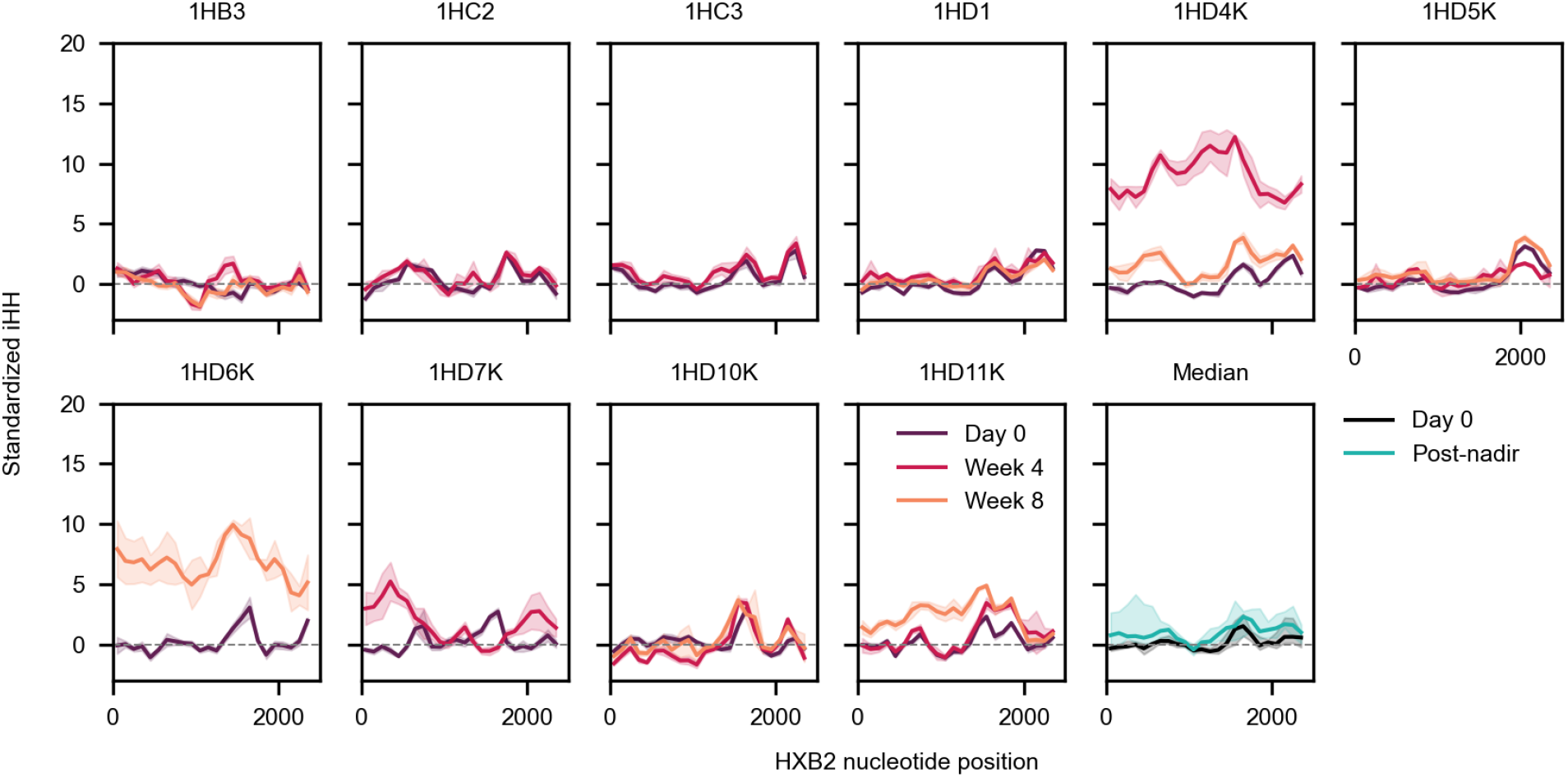
Haplotype length did not change in most HIV populations following 10-1074 infusion. Standardized iHH is plotted versus HXB2 nucleotide coordinate at different time points preceding and following 10-1074 infusion (Materials & Methods). Standardized iHH is plotted as a binned average (100 nucleotide positions per bin) with shading showing 95% confidence intervals on each bin. The final panel shows the median standardized iHH across all participants in each genomic bin. These median values are shown at day 0, prior to infusion, and at the first time point post viral load nadir, with the IQR indicated via shading.

Following the spread of adaptive alleles, we observed an increase in iHH in participants 1HD4K and 1HD6K characteristic of selective sweeps. In 1HD4K, standardized iHH increased from a mean of approximately zero to 9.04 *±* 3.46 across Env at week 4, corresponding with the timing of viral load reaching pre-treatment levels (Figure 2A). By week 8, iHH returned to near pre-sweep levels, reflecting the transiency of haplotype signatures in this setting. Similarly, participant 1HD6K had an elevation of iHH at week 8 corresponding to the first timepoint at which their viral load was above 10^3^ copies/mL. Note, no viral sequences from week 4 are available for 1HD6K. Although 1HD4K experienced a modest reduction in diversity as described above, 1HD6K’s iHH patterns confirm that haplotype signatures can capture soft sweeps when diversity signatures cannot.

Remarkably, however, in the majority of participants (8/10), we found negligible change in haplotype length four weeks after therapy (Figure 4). Despite the rapid emergence of escape mutations in all 10-1074 treated intra-host HIV populations, iHH values often overlapped pre-treatment levels. Further, standardized iHH curves in post-rebound viral populations often recapitulated slight regional variations in standardized iHH at pre-treatment time points, indicating haplotype features are preserved during adaptation (Figure 4). These patterns suggest that HIV can escape 10-1074 via sufficiently soft sweeps that resistant populations show no clear signals in iHH.

We term sweeps that minimally affect diversity or haplotype length as “ultra-soft” selective sweeps.

### Simulated selective sweeps contextualize how iHH and diversity signatures change as sweeps become softer

Having identified that adaptation during 10-1074 escape does not exhibit traditional genetic sweep signatures, we next used simulations to characterize how the genetic signatures of selective sweeps change with increasing sweep softness.

To contextualize our observations in the 10-1074 dataset, we simulated matched evolutionary conditions to the 10-1074 clinical trial. We simulated haploid populations of constant size using SLiM [Haller et al., 2025]. We parametrized the simulations using existing HIV estimates for the effective population size [Achaz et al., 2004], mutation rate [Zanini et al., 2017], distribution of fitness effects [Zanini et al., 2017], and recombination rate [Romero and Feder, 2024] (See Materials & Methods). These simulations produced full *env* length nucleotide sequences, which were sampled to roughly match the average sequencing depth per time point in Romero et al. [2026].

We first tested that our simulations matched the *in vivo* pre-treatment nucleotide diversity, the site frequency spectrum, and *D^′^* as a function of genetic distance within participants. We found that our pre-sweep simulated populations had comparable values in terms of nucleotide diversity (Figure S3A), the relative number of rare and common alleles (Figure S3B), and linkage decay rate (Figure S3C). There were individual participants in which one or more metrics fell away from the simulated mean (for example, we found fewer rare alleles in participant 1HB3), but we concluded that our simulated conditions were a plausible baseline for understanding post-sweep diversity. We therefore proceeded with simulating selective sweeps in these diversity-matched populations to profile their corresponding genetic signatures when different numbers of adaptive mutation origins drive escape.

We introduced between 1 and 100 “sweep origins” by randomly sampling sequences and placing highly beneficial mutations onto each of them at a single escape locus (*s* = 3.33, Materials & Methods). Then we monitored the sweeping populations over the course of 22 generations to roughly match the time course of the existing 10-1074 trial data [Perelson et al., 1996, Caskey et al., 2017]. We retained only simulations where a sufficient number of sweep origins remained present throughout the sweep (Figure S4, See Materials & Methods).

Tracking diversity signatures throughout these simulated trials, we first confirmed that single origin hard sweeps depressed genetic diversity surrounding the selected locus [Maynard and Haigh, 2007]. Strong selection and a short genic length resulted in depressed diversity nearly across the full length of *env* (Figure 5). As the number of sweep origins increased (2-10 origins), diversity loss (as measured via change in average pairwise Hamming distance) became less extreme, more variable across replicates, and was only detectable when aggregating across multiple simulations. When more than 10 origins were simulated, we detected no consistent diversity loss in the region surrounding the selected locus. Mirroring our observations in the *in vivo* dataset, diversity actually increased during the trial at the escape locus, driven by the multiple distinct beneficial alleles simultaneously expanding. Furthermore, when we simulate soft sweeps of beneficial mutations spread across different escape regions, we observe these diversity increases at all of the escape loci (Figure S5). Consistent with other studies [Feder et al., 2016, Anderson et al., 2017], our results suggest that soft sweeps can permit adaptation with minimal discernible changes in diversity in the region surrounding a selected locus and increases in diversity at the selected locus itself.

**Figure 5:**
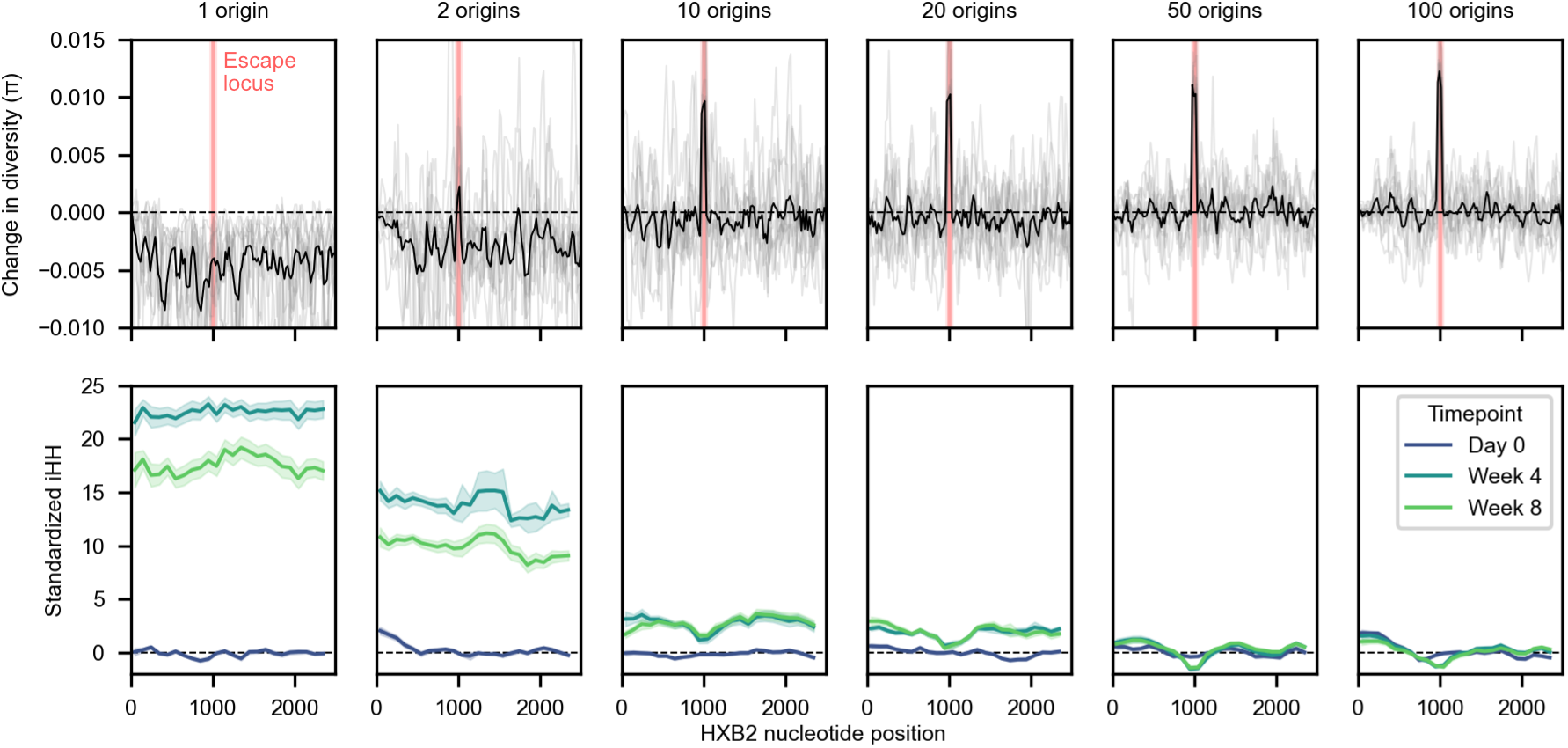
Classical diversity and haplotype based sweep signatures lessen as the number of origins grow. **(Top)** Change in average pairwise Hamming distance between trial “day 0” and trial “week 4” (11 simulated generations later) for sweeps originating from 1 to 100 highly beneficial mutations (*s* = 3.33) introduced into randomly selected HIV genomes at a given escape locus (Materials & Methods). Individual simulation replicates are shown in light gray while the mean over 10 replicates is shown in black. Hamming distance is calculated in 60 nucleotide bins advanced by 20 nucleotides. The escape locus is shaded red. **(Bottom)** Standardized integrated haplotype homozygosity (iHH) plotted versus HXB2 nucleotide coordinate during simulated selective sweeps of different numbers of origins (Materials & Methods). Data are pooled across 10 replicates and plotted as a binned average (100 nucleotide positions per bin) with 95% confidence intervals on each bin.

We next quantified the impact of sweep softness on haplotype signatures using standardized integrated haplotype homozygosity (iHH). At one or two sweep origins, iHH throughout the genic region sharply increased directly following the selective sweep (i.e., week 4). In subsequent post-sweep generations (i.e., week 8), iHH remained elevated but decreased towards the pre-sweep baseline, similar to the patterns observed in participant 1HD4K. iHH was also modestly elevated post-sweep for larger numbers of sweep origins. In sweeps of 10 and 20 origins, week 4 and 8 standardized iHH remained elevated above the day 0 baseline, both on average (Figure S5) and in individual simulations (Figure S6). Haplotype lengths therefore indeed show more robust signatures of sweeps across a wider parameter range than relying on diversity alone.

However, as the number of origins approached and exceeded 50, simulated populations showed no elevation in iHH following selective sweeps, similar to certain participants within the 10-1074 trial dataset. While single simulation replicates are noisy, they show no consistent temporal signal in iHH with respect to the sweep (Figure S6). We did observe a reduction in iHH localized to the escape locus at higher numbers of origins (Figures 5, S5). This effect is driven by single mutations at the escape locus itself linked to multiple genetic backgrounds. However, both the *in vivo* and single replicate data appear to be too noisy to reliably detect this signal (Figures 4, S6). Because the haplotype structure of pre and post-sweep populations is largely identical at higher numbers of origins, these ultra-soft sweeps are essentially indistinguishable via iHH from scenarios with no adaptation.

Our simulations reveal that classical signatures of selective sweeps that are apparent at few origins can completely disappear and, in some cases, invert when sweeps are instead driven by large numbers of origins. As expected, linkage statistics better capture the perturbation of softer selective sweeps than diversity statistics do, but both ultimately fail when pushed further into the ultra-soft regime.

### *In vivo* iHH is consistent with 20 or more escape origins

To summarize the haplotype signature results from simulations and compare them to the *in vivo* data more directly, we computed a new statistic to measure change in iHH following a sweep. Our approach parallels the integrated haplotype score (iHS), in which iHH among haplotypes carrying a derived allele is compared to iHH for those carrying the ancestral allele [Voight et al., 2006]. Because we have longitudinal data available, we can directly compare pre- and post-sweep iHH patterns instead. We report the signed area between the iHH curves at pre- and post-sweep time points, and refer to the resultant statistic as integrated iHH (iiHH, see Materials and Methods). Consistent with direct examination of the iHH curves, measurable elevations in post-sweep haplotype length are apparent. Simulations with 20 or fewer sweep origins result in positive iiHH values, while the minimal perturbations in post-sweep iHH with 50 or more origins result in iiHH distributions tightly centered near 0, indicating no change to the length of haplotypes following a sweep (Figure 6A). We found that most of the 10-1074 trial participants had low iiHH values, consistent with higher origin number sweeps. Two participants (1HD6K and 1HD4K) had higher iiHH values.

**Figure 6:**
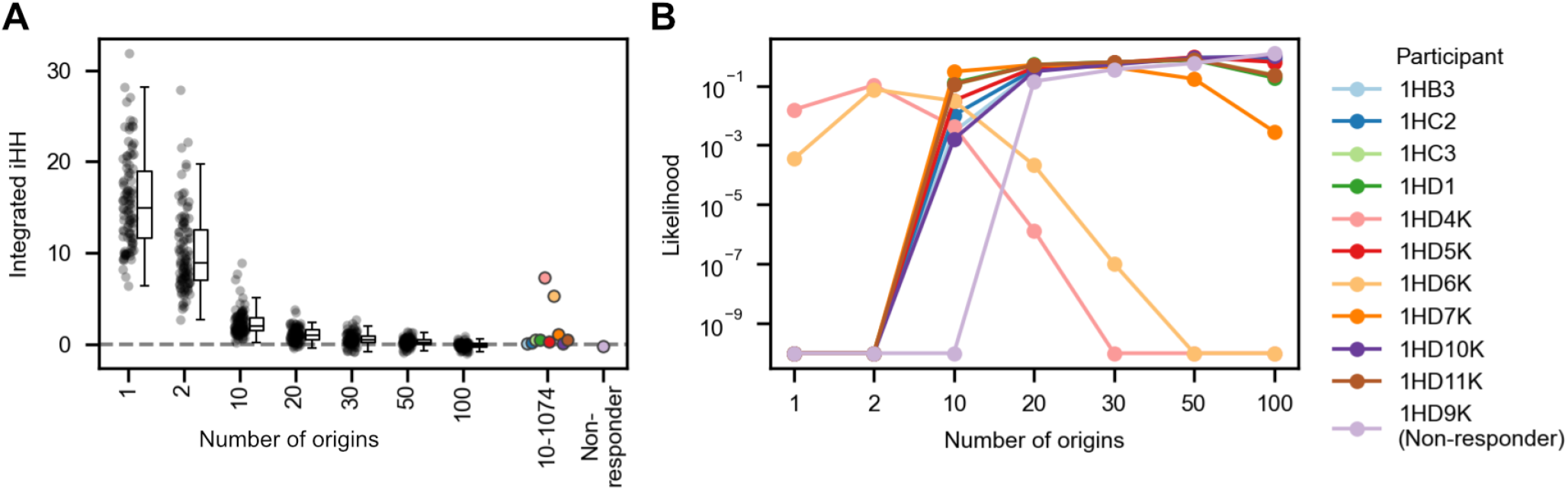
Integrated iHH (iiHH) during 10-1074 escape is consistent with 20 or more sweep origins in most participants. **(A)** iiHH values for simulations with different numbers of sweep origins are shown alongside the iiHH values for the 10 viral populations escaping 10-1074 and the one non-responding viral population. For each number of simulated origins, a strip plot shows iiHH for 100 replicates and a boxplot summarizes over all replicates. iiHH is calculated as the signed area between standardized iHH curves (Figure 4) before and after escape (Materials & Methods). **(B)** Likelihood of the number of sweep origins for each participant given their iiHH value.

We next used these distributions of iiHH to roughly estimate the number of sweep origins in each participant’s escaping intra-host HIV population. We fit each distribution of iiHH resulting from a specific range of simulated sweep origins to a gamma distribution and computed the likelihood of participant-specific iiHH values under each distribution (see Materials and Methods, Supplemental Figure S7). We found that the high values of iiHH in 1HD6K and 1HD4K were consistent with a relatively small number of origins, while iiHH values in nearly all other participants were better explained by 20 or more sweep origins (Figures 6B). These estimates are consistent with the lower bounds we previously estimated by tracking haplotypes carrying escape mutations [Romero et al., 2026]. Although we did not simulate exhaustively across the entire range of possible sweep origins, the saturation of these likelihood curves at greater numbers of sweep origins makes clear that we have limited resolution to distinguish among ultra-soft sweeps using iiHH.

As a useful point of comparison, the 10-1074 cohort also happened to contain an additional participant (1HD9K) who did not respond to therapy (Figure S8A). Their viral population was later found to be fixed for a pre-existing escape mutation before the trial began, which was confirmed phenotypically as unresponsive to 10-1074 *in vitro* [Caskey et al., 2017]. As a result, they did not experience a change at any of the escape loci during the trial period. Notably, this participant had an iiHH value of −0.14 at week 4 of the trial, which denotes a slight iHH decrease and is similar to those HIV populations experiencing ultra-soft sweeps, despite 1HD9K’s lack of sweep. Likelihood fits to their iiHH value revealed it to be most consistent with the highest numbers of origins in sweeps that we simulated (80-100), and their iHH signatures were otherwise similar to other participants who experienced resistance evolution during the trial period (Figure S8).

Collectively, these comparisons show that the minimal haplotype based signatures of ultra-soft selective sweeps can, to some extent, be used to estimate the number of origins underlying already known sweeps, such as those identified by neutralization assays and viral load rebound in the 10-1074 clinical trial data. However, detecting adaption via ultra-soft sweeps outside of known circumstances, like the clinical trial studied here, may be challenging from iHH signatures alone because those signatures can be highly similar to non-response.

## Discussion

Widespread genome sequencing has revealed that many canonical instances of adaptation are in fact driven by soft sweeps derived from multiple mutational events. Here, we demonstrate that when HIV evolves to escape 10-1074 monotherapy, it often does so with sweeps so soft as to leave effectively no impact on either the genetic diversity or haplotype length of the neighboring genomic regions. Comparison to simulations suggests that, for the majority of trial participants, escape mutations likely exist on 20 or more intra-host genetic backgrounds, although further differentiating between the softnesses of these ultra-soft sweeps is challenging due to their convergence to minimal genetic signatures at higher numbers of origins in this system.

The extreme softness of these sweeps has clinical implications for HIV infections and their treatment. Recent trials are increasingly combining broadly neutralizing antibodies such as 10-1074 to slow resistance evolution [Julg et al., 2024, Sneller et al., 2022]. Quantifying the number of origins of adaptive alleles in monotherapy escape, even coarsely, informs how many therapies would be necessary to combine to fully limit escape [LaMont et al., 2022]. The widespread maintenance of haplotypes and individual mutations following these ultra-soft sweeps also has potential implications for the onward evolutionary trajectories of these viral populations. Immunological control of HIV populations is enhanced when viruses have limited opportunity to diversify [Hocqueloux et al., 2010]. Therefore, acquiring resistance while retaining nearly all intra-host HIV population diversity may be challenging for the host across multiple axes. Relatedly, we previously found that the simultaneous availability of multiple 10-1074 escape mutations permitted the virus to select those that minimize antagonistic pleiotropy for other important viral functions like cell entry [Romero et al., 2026]. More broadly, the ultra-soft sweeps described here may have a previously unrecognized role in HIV escape from autologous antibodies, although this remains to be demonstrated.

Although most of the HIV populations analyzed here showed iiHH patterns consistent with *≥* 20 origins of resistance, 1HD4K and 1HD6K did not. We do not have a clear explanation for why these populations might be different. Theory predicts that these relatively hard sweeping populations might start with smaller population sizes and less genetic diversity. While 1HD4K and 1HD6K both had lower than average pre-treatment diversity within the cohort, they were not outliers, and we found no evidence for a relationship between sweep hardness and viral load (Figure S9). Notably, participant 1HD6K had the longest time to viral load rebound following 10-1074 infusion (Figure 2), which is consistent with a lower mutational input and resultant harder selective sweep. However, it is unclear why 1HD6K’s mutational input would be lower based on pre-treatment characteristics. In contrast, participant 1HD4K experienced immediate rebound with the smallest transient reduction in viral load within the cohort (Figure 2). At week 4, 69% of 1HD4K’s sequences carried N332D, a mutation observed only at low frequencies in a subset of the other participants. No other participant had a single resistance mutation at this high of a frequency. One explanation for the modest reduction in viral load and prevalence of N332D is that this mutation pre-existed at unexpectedly high frequency before the onset of therapy, which permitted the population a fast rebound driven by N332D before other variants could establish. However, we did not detect N332D at the pre-infusion sampling time, while we did detect a single copy of S334N. Why these HIV populations behave differently therefore remains an open question.

We are not the first to report instances of very soft selective sweeps leaving little impact on diversity [Kreiner et al., 2019, Cao et al., 2025, Anderson et al., 2017, Warwick et al., 2008, Feder et al., 2016]. However, to our knowledge, this report is one of a very small number that demonstrate soft selective sweeps without clear alterations in the haplotype structures of the populations. One such example is Laurin & Garud 2026, who found widespread adaptation at the *gyrA* locus which is often altered in antibiotic resistance. Across single bacterial sequences across many distinct hosts, the linkage statistic H12 [Garud et al., 2015] is not elevated within *gyrA*, despite widespread phenotypic resistance [Laurin and Garud, 2026]. The authors suggested the explanation that these single nucleotide variants were produced independently in each population without spreading across hosts. Kreiner et al. also found no perturbation of haplotype structure measured by XP-EHH [Sabeti et al., 2007] among certain populations of glyphosate-resistant amaranthus [Kreiner et al., 2019, 2022]. It is possible this pattern is present in other settings, but many studies report only variants associated with resistance without clear analyses of associated haplotypes.

Why then has this limited haplotype signature been rarely observed previously? One possibility relates to how these populations are sampled. Many studies of adaptation compare samples across multiple geographic regions where sweeps may appear locally hard despite being globally soft. In contrast, our investigation worked with plasma samples of HIV virions pooled from many anatomical locations, which may obscure locally hard signals otherwise revealed by spatial sampling. For example, Feder et al previously observed partially distinct sweeps of drug resistance mutations in gut, lymphoid and vaginal tissues of drug-treated SHIV-infected macaques [Feder et al., 2017]. These mutations were all present in the plasma, making the sweep appear softer than in any individual compartment. Similar patterns of locally hard, globally soft sweeps are also observed in other populations [Cao et al., 2025, Délye et al., 2013, Nair et al., 2007]. However, such patterns can be obscured even in regionally-sampled data with sufficient gene flow [Kreiner et al., 2019, 2022]. Therefore, while the extremely subtle signatures of the ultra-soft sweep described here may be mitigated when sequencing spatially distinct samples, understanding the associated signatures of ultra-soft sweeps pooled from many regions remains important.

It is important to note that these particular sweeps were indeed visible despite the limited diversity and iHH signatures; three loci showed substantial genetic divergence between pre and post-treatment populations. Similar attributes have been central to identifying very soft sweeps of resistance to artemisinin, ALS–inhibitors, glyphosate, and other biocides [Anderson et al., 2017, Warwick et al., 2008, Kreiner et al., 2019, Cao et al., 2025, Nair et al., 2007]. However, observing repeated changes at few sites may not be possible if the mutational target size is large and/or distributed across the genome such that the signature at any given locus is small. For example, we previously analyzed HIV populations treated with a different broadly neutralizing antibody, 3BNC117, from which escape can occur at many sites Romero et al. [2026], Radford and Bloom [2025]. In some of these populations, distinguishing between no sweep and an ultra-soft sweep was difficult in the absence of phenotypic data. Therefore, it remains necessary to develop robust approaches to detect ultra-soft sweeps in the absence of the most widely-appreciated haplotype and diversity signals.

We note several promising developments. First, methods that detect allele frequency changes from time series data have successfully identified even extremely soft sweeps concentrated at few loci [Sohail et al., 2021, Romero et al., 2026]. However, the utility of these tools may become more limited when more mutations spread if clonal interference limits the spread of any individual mutation [Lang et al., 2013, Gerrish and Lenski, 1998]. Second, the subtle post-sweep signatures observed here parallel those generated by polygenic adaptation in which many alleles of small effect shift slightly in frequency [Pritchard et al., 2010, Barghi et al., 2020, Höllinger et al., 2019]. Emerging tools in this field may have some transferability to discover ultra-soft sweeps. We attempted to apply one such statistic (the Singleton Density Score) and found we had too few singletons within our dataset [Field et al., 2016]. However, approaches to detect distortion in rare (as opposed to common) variation may be portable. Third, advances in high throughput screening techniques allow rapid phenotyping of populations to detect population changes even without clear genetic signatures. Prospective mapping of mutational effects via techniques like deep mutational scanning could enable scans that look for those alleles directly [Radford and Bloom, 2025, Dingens et al., 2019, Pendyala et al., 2026]. Finally, we note that haplotype-based approaches like iiHH retain evidence of soft selective sweeps on as many as 20 origins of resistance in HIV-like simulation conditions, which covers many of the cases of even very soft sweeps in the published literature.

We note some caveats and future directions. First, we focused our investigation on sweeps stemming primarily from a single adaptive locus under HIV-like evolutionary conditions. Although we found similar results when examining the impact of more dispersed adaptive loci, additional study should further probe the generalizability of our results. For example, since diversity and haplotype based detection methods rely on segregating background variation at flanking sites, strong purifying selection or low mutation rates may further obscure ultra-soft sweep signatures. Second, we considered a simplified case in which adaptive alleles were introduced at the onset of therapy instead of segregating in populations ahead of therapy. We therefore cannot distinguish scenarios where adaptive alleles arose from multiple mutational origins and scenarios where a single adaptive allele arose via mutation and then recombined onto different haplotypes. We chose this approach because 10-1074 escape alleles are associated with strong fitness costs and are constrained to very low frequencies in the absence of therapy [Radford and Bloom, 2025, Romero et al., 2026, Caskey et al., 2017]. However, we urge caution in interpreting the number of origins as unique mutational events instead of the number of genetic backgrounds upon which adaptive alleles spread. Third, our simplified model encoded only three distinct nucleotides as escape mutations while HIV escaped from 10-1074 via a median of *≈*9 (range: 4 to 15) different nucleotide substitutions in Romero et al. [2026]. This discrepancy reduced the height of the diversity peak at the escape locus and magnitude of the iHH reduction surrounding the escape locus in simulations, which depend on the exact nucleotide identities of escape mutations. However, the multi-locus simulations had higher numbers of nucleotide identities causing escape but retained similar characteristics (Figure S5).

In summary, we use HIV escape from the broadly neutralizing antibody 10-1074 to characterize a mode of adaptation driven by many independent mutational origins, which we term an “ultra-soft” selective sweep. We show that as sweeps become progressively softer, their signatures of haplotype homozygosity and genetic diversity become so weak that they become indistinguishable from scenarios without any adaptation at all. This may limit our ability to identify other ultra-soft sweeps in genomic data. Because they preserve standing genetic variation and haplotype diversity, ultra-soft sweeps may enable populations to repeatedly adapt to new or concurrent selective pressures without sacrificing future adaptive potential.

## Materials & Methods

### *In vivo* Data

The samples profiled in this paper were originally collected as part of NCT 02511990, a clinical trial that administered a single infusion of broadly neutralizing antibody 10-1074 to viremic people living with HIV [Caskey et al., 2017]. HIV from plasma samples collected longitudinally at therapy initiation (day 0), week 1, week 4 and week 8 was sequenced to approximately 20*×* depth using single genome sequencing [Salazar-Gonzalez et al., 2008]. A subset of trial participants (1HB3, 1HC2, 1HC3, 1HD1, 1HD4K, 1HD5K, 1HD6K, 1HD7K, 1HD9K, 1HD10K, and 1HD11K) were deeply resequenced (median 40 sequences per sample; range: 1 to 198) in Romero et al. [2026] using SMRT-UMI sequencing [Westfall et al., 2024]. We profiled viral populations in all participants with deep sequencing data except for participant 1HD9K, whose viral population carried a fixed pre-existing escape mutation (D325E) and did not respond to 10-1074. Sequences from both the original study [Caskey et al., 2017] and the re-sequencing study [Romero et al., 2026] are included in our analysis. Supplemental figure S10 shows all sequencing time points analyzed here. Sequences from each participant were quality filtered and aligned as described in Romero et al. [2026], and the full list of analyzed sequences is listed in Supplemental Table 3 in that paper.

All sequences with either mutations removing the N332 potential N-linked glycosylation site or mutations away from D/N325 were considered to have escaped 10-1074 [Caskey et al., 2017, Dingens et al., 2019, Radford and Bloom, 2025].

### Simulated Data

We performed forward simulations in SLiM version 5.1 [Haller et al., 2025]. We used the nucleotide-based model to simulate the full HIV *env* gene (2571 nucleotides) initialized to the HXB2 reference sequence. We simulated haploid populations with a mutation rate of *µ* = 10*^−^*^5^ [Achaz et al., 2004, Zanini et al., 2017, Sanjúan et al., 2010], recombination rate of *ρ* = 7.6 *×* 10*^−^*^6^ [Romero and Feder, 2024, Neher and Leitner, 2010, Longley et al., 2025], and constant effective population size *N_e_*= 10^4^ [Feder et al., 2017, Achaz et al., 2004]. We simulated constant population sizes rather than more complex demographic models because previous investigations observed only modest differences in sweep signatures between constant size populations and those rescued from extinction by adaptive mutations [Osmond and Coop, 2020]. New mutations were deleterious (*s* = *−*0.1), neutral (*s* = 0) or beneficial (*s* = 0.05) with probabilities 0.665, 0.333 and 0.001, respectively, reflecting Zanini et al. [2017]’s finding that most nonsynonymous mutations have a *>* 10% fitness cost and most synonymous mutations have a *<* 1% fitness cost. 100 genomes were sampled per time point to roughly match the average sampling depth of the combined *in vivo* data (70 sequences per time point). Generations were converted to approximate time points using an approximate generation time of 2 days per generation [Perelson et al., 1996].

#### Initial HIV parameter matching

We first used simulations without artificially-introduced selective sweeps to verify the simulated scenarios matched the pre-trial *in vivo* sequence data. These simulations each ran for 500 generations and were sampled every 50 generations. We first matched simulations based on average pairwise Hamming distance and found that generation 300 best matched the standing genetic diversity of the *in vivo* populations before 10-1074 infusion. Using the generation 300 time point, we visually verified that these simulated parameter settings also matched the relative balance of rare and common alleles in the folded allele frequency spectrum, and the decay of linkage as a function of distance between two loci as measured by D’.

#### HIV selective sweep simulations driven by variable numbers of adaptive alleles

We ran simulations for 300 generations to generate comparable diversity, linkage and site frequency spectra to the pre-infusion *in vivo* data (see above). We then artificially introduced 1, 2, 10, 20, 30, 50, or 100 strongly beneficial alleles randomly on to pre-existing genetic backgrounds. We sampled the population at generations 300, 303, 311 and 322 to match the trial sampling scheme of 0, 1, 4, & 8 weeks. If fewer than a given number of adaptive alleles are circulating at any time between generations 300 and 322 (1, 2, 5, 10, 20, 40, or 80, respectively) or if adaptive alleles do not reach 99.9% of the population by generation 322, the simulation is restarted at generation 300. Distributions of surviving origin numbers are shown in Figure S4.

Artificially-introduced adaptive alleles were given a selection coefficient of *s* = 3.33 to permit a full sweep in 22 generations. Unless otherwise noted, they were always introduced at nucleotide 1000 to align with the *in vivo* escape positioning and so that each sequence contained only a single escape mutation. Escape mutations were any of the three nucleotide substitutions away from the ancestral allele, although each distinct mutational event is separately tracked via its own ID in SLiM to allow conditioning on sufficient mutational survival. Note, this is a simplified encoding of the escape locus architecture for 10-1074 (see Discussion).

We also relaxed the assumption of all mutations occurring at only a single nucleotide position. We performed additional simulations in which escape alleles were randomly added anywhere in HXB2 nucleotide coordinates [808-855], [1078-1113], and [1363-1455], mirroring the contact sites for a different broadly neutralizing antibody, 3BNC117 [Schoofs et al., 2016, Zhou et al., 2013]. Escape loci and nucleotides changes were drawn randomly for each origin, and sequences were limited to carrying only a single escape mutation via SLiM’s modifyChild() callback [Haller et al., 2025].

### Summary statistics

#### Average pairwise diversity

For each sample and time point, we calculated nucleotide diversity using average pairwise Hamming distance in a 60 nucleotide sliding window advanced by 20 nucleotides across the genome. For each pair of focal sequences, positions where either sequence had a gap were filtered out prior to calculating their differences.

#### D’

To match linkage disequilibrium as a function of distance separating two alleles between our empirical and simulated data, we computed D’ between pairs of alleles [Lewontin, 1988]. For each sample and time point, segregating sites with minor allele frequencies *<* 5% were filtered out, then D’ was calculated for all pairs of remaining segregating sites.

#### Integrated haplotype homozygosity (iHH)

We quantified haplotype length using integrated haplotype homozygosity (iHH), a measure derived from extended haplotype homozygosity (EHH) [Sabeti et al., 2002, Voight et al., 2006]. Note, we used iHH for the majority of our analyses due to its greater sensitivity for detecting linkage changes than *D^′^*, *r*^2^ and other related linkage statistics in preliminary analyses. For each allele at every segregating site, we calculated EHH as the probability that two randomly selected sequences carrying the focal allele are identical at all sites between the focal locus and a given position Sabeti et al. [2002]. We calculated EHH at increasing distances from the focal locus to generate an EHH decay curve and integrated this curve to obtain iHH [Voight et al., 2006]. Analyses were restricted to segregating alleles with a frequency greater than 0.05 and at least two observations across different sequences.

To account for the dependence of iHH on allele frequency, we standardized iHH values following Voight et al. [2006]:

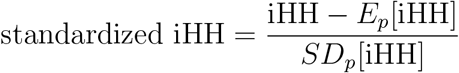

where *E_p_*[iHH] and *SD_p_*[iHH] are the mean and standard deviation of iHH among alleles with frequency *p*. These quantities were estimated from the empirical distribution of alleles in the day 0 sample for the corresponding individual or simulation. Allele frequencies were grouped into bins of width 0.2 to ensure sufficient coverage.

#### Integrated integrated haplotype homozygosity (iiHH)

To characterize gene-wide changes in iHH over time, we calculated standardized iHH values for all segregating loci and averaged them across 250 equally spaced genomic bins (approximately 10 nucleotides per bin). This process produced averaged iHH curves, where each point corresponded to the bin midpoint and the average iHH within that bin, with adjacent points connected by linear interpolation. To quantify changes in iHH between two timepoints, we calculated the signed area between their corresponding curves. Any bins with no data in either comparison curve were dropped, and linear interpolation was instead performed between the flanking bins on either side. Positive integrated iHH values indicate an overall increase in haplotype lengths relative to day 0, whereas negative values indicate a decrease.

Because both our empirical data and simulations are computed over the same sequencing length (2571 nucleotides), we divided all iiHH values by 2571 for ease of visualization. However, we caution that iiHH should actually be approximately independent of the sequenced length if EHH declines back to 0. Therefore, if comparing iiHH values between two different scenarios with different sequenced lengths (i.e., chromosomes of different lengths), this standardization should not be applied.

### Likelihood analysis

We computed the likelihood that each of our *in vivo* iiHH values was drawn from a distribution of iiHH values in simulations with a given origin. For each origin number in the simulated data, we fit a gamma distribution to the resulting iiHH values for that set of simulations using the scipy.stats.gamma.fit() function (Figure S7, Table S1) [Virtanen et al., 2020]. Then, we calculated the likelihood that the *in vivo* iiHH value for each participant was drawn from each gamma distribution.

### Allele Persistence Analysis

We tracked all segregating alleles that were present at day 0 and reached frequency *>* 5% during at least one trial time point. We defined alleles as persisting if they were observed during at least one trial time point post viral load nadir in the corresponding study participant.

We then formulated a null expectation of how many alleles would persist if the observed allelic loss was due to sequencing depth alone. For each segregating site at each time point *in vivo*, we drew a random allele frequency from the participant’s distribution of allele frequencies across all time points. Then, we drew from a binomial distribution with success probability *p* set to the drawn frequency and number of trials *n* set to the number of sequences collected from the participant at the given time point. We marked the allele as sampled if it was drawn at least once. We repeated this process 1000 times for each participant to form a bootstrapped null dataset.

We separately trained logistic regression models on the null and *in vivo* datasets using statsmod-els fit regularized() [Seabold and Perktold, 2010]. These models were of the form

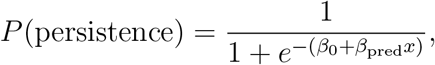

where *β*_pred_ was the predictor variable. We fit different regression models using either allele frequency or distance from escape locus as predictor variables.

## Acknowledgements

We would like to thank the Feder lab members, Nandita Garud, Kelley Harris, Sophie Seidel and Pleuni Pennings for thoughtful feedback on this manuscript.

## Code Availability

All code used for conducting analyses and producing figures for this work is available at https://doi.org/10.5281/zenodo.21465309.

## Data Availability

All 10-1074 trial sequences from Romero et al. [2026] are available in GenBank under accession numbers PX290569-PX292892 while all sequences from Caskey et al. [2017] are available under accession numbers KY323724–KY324834.

## Supplement

**Figure S1:**
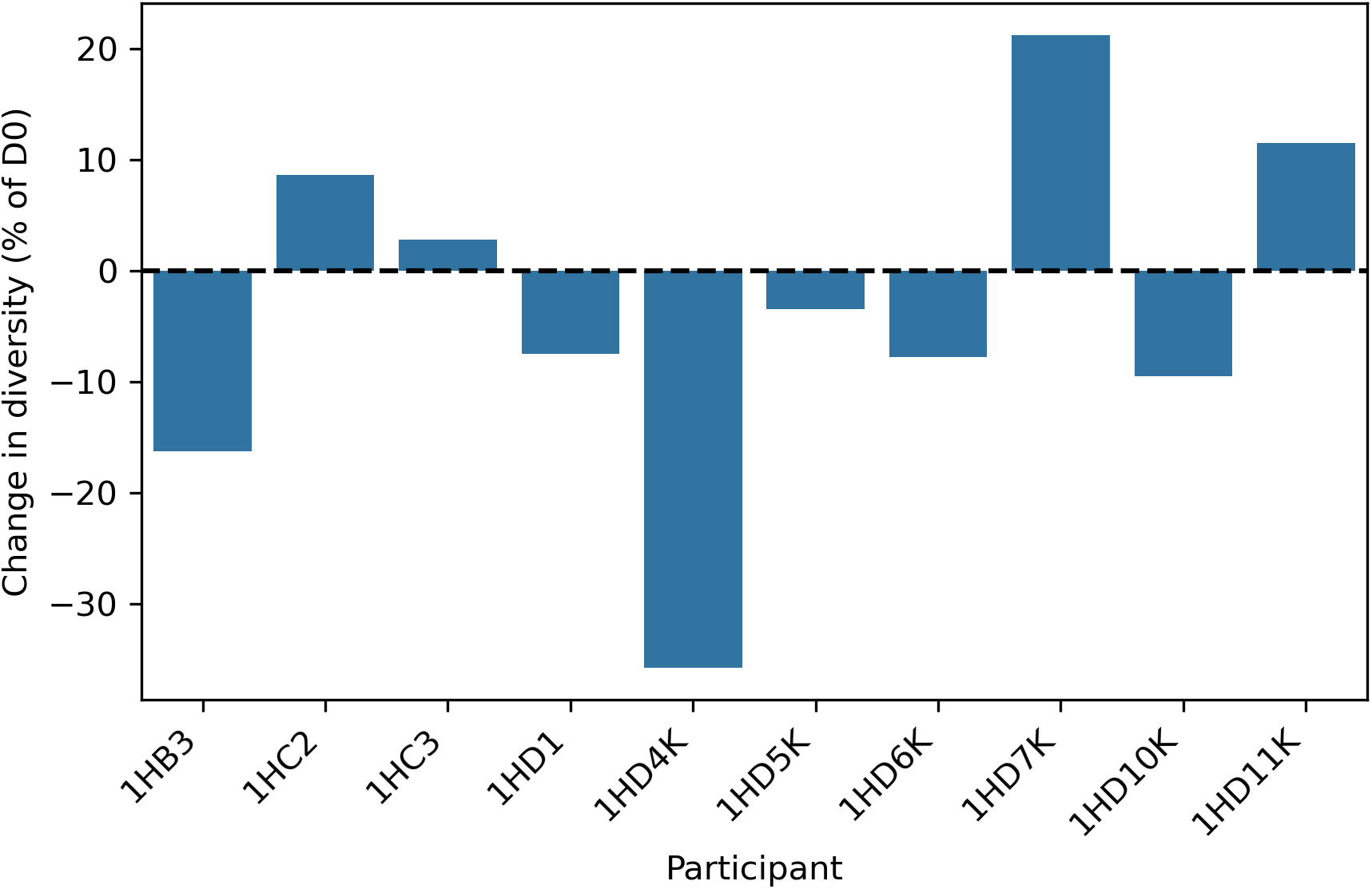
Change in average pairwise Hamming distance across *env* following 10-1074 infusion. Change in average pairwise Hamming distance across the full *env* gene between trial day and the first time point following viral load nadir is shown for each trial participant.

**Figure S2:**
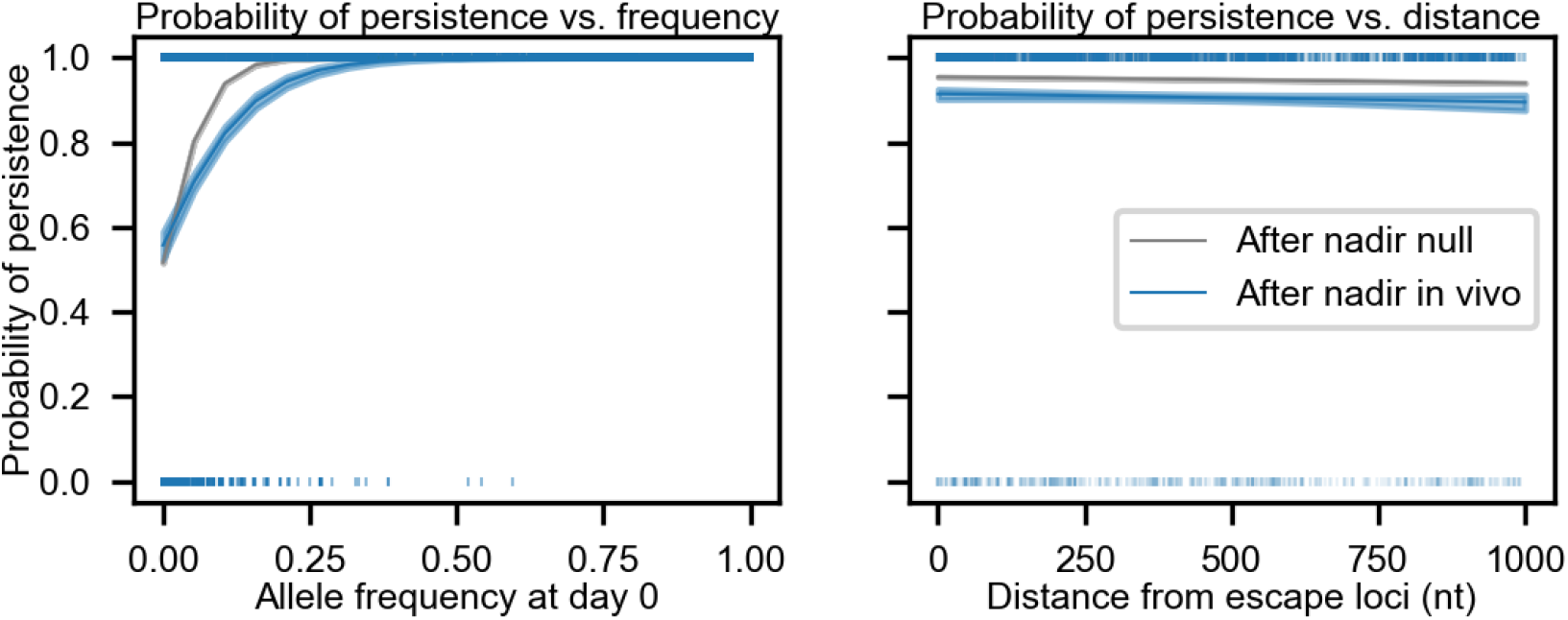
Mutational probability of persistence following a sweep depends on pre-sweep frequency but not distance from the selected loci. Tick marks at *y* = 1 or *y* = 0 indicate alleles segregating at day 0 and if they persisted or were lost, respectively, during the trial. The x-axis plots their day 0 allele frequency within a participant (left) or distance from the escape loci (right). Data from all participants is shown. Logistic regression fits to these data are plotted in blue. Null expectations of allelic loss based on the depth of sequencing at each time point are shown in gray (See Materials & Methods).

**Table S1:**
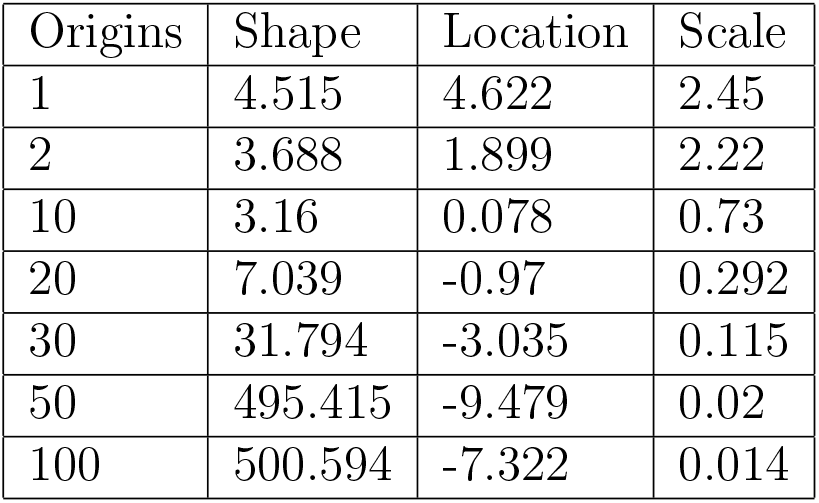
Parameters for gamma distributions fit to the simulated iiHH distributions that are shown in figure S7.

| Origins | Shape | Location | Scale |
| --- | --- | --- | --- |
| 1 | 4.515 | 4.622 | 2.45 |
| 2 | 3.688 | 1.899 | 2.22 |
| 10 | 3.16 | 0.078 | 0.73 |
| 20 | 7.039 | -0.97 | 0.292 |
| 30 | 31.794 | -3.035 | 0.115 |
| 50 | 495.415 | -9.479 | 0.02 |
| 100 | 500.594 | -7.322 | 0.014 |

**Figure S3:**
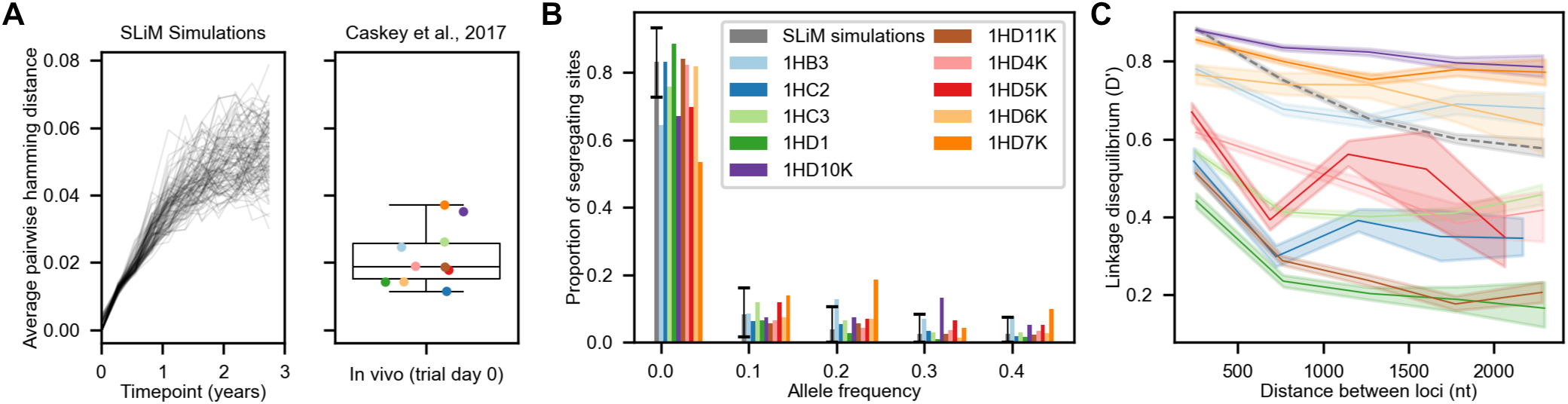
Simulations match *in vivo* pre-treatment diversity, allele frequency spectra, and linkage decay. **(A)** Left: Average pairwise Hamming distance over time in 100 simulated intra-host HIV populations. Right: Average pairwise Hamming distance in trial participants at trial day 0, before 10-1074 infusion [Caskey et al., 2017, Romero et al., 2026]. **(B)** Minor allele frequency spectrum for simulated intra-host populations (gray) and for 10-1074 trial participants at day 0 (colored). Simulated populations are sampled at generation 300 and error bars indicate 95% confidence intervals across 100 replicate simulations. Spectra are binned in increments of 0.1. **(C)** Relationship between *D^′^*[Lewontin, 1988] and separating distance between pairs of segregating loci. The gray dashed line indicates the median linkage decay in 100 replicate simulations while the colored lines indicate the median linkage decay within each study participant. Shading represents 95% bootstrapped confidence intervals (1000 bootstrapped replicates).

**Figure S4:**
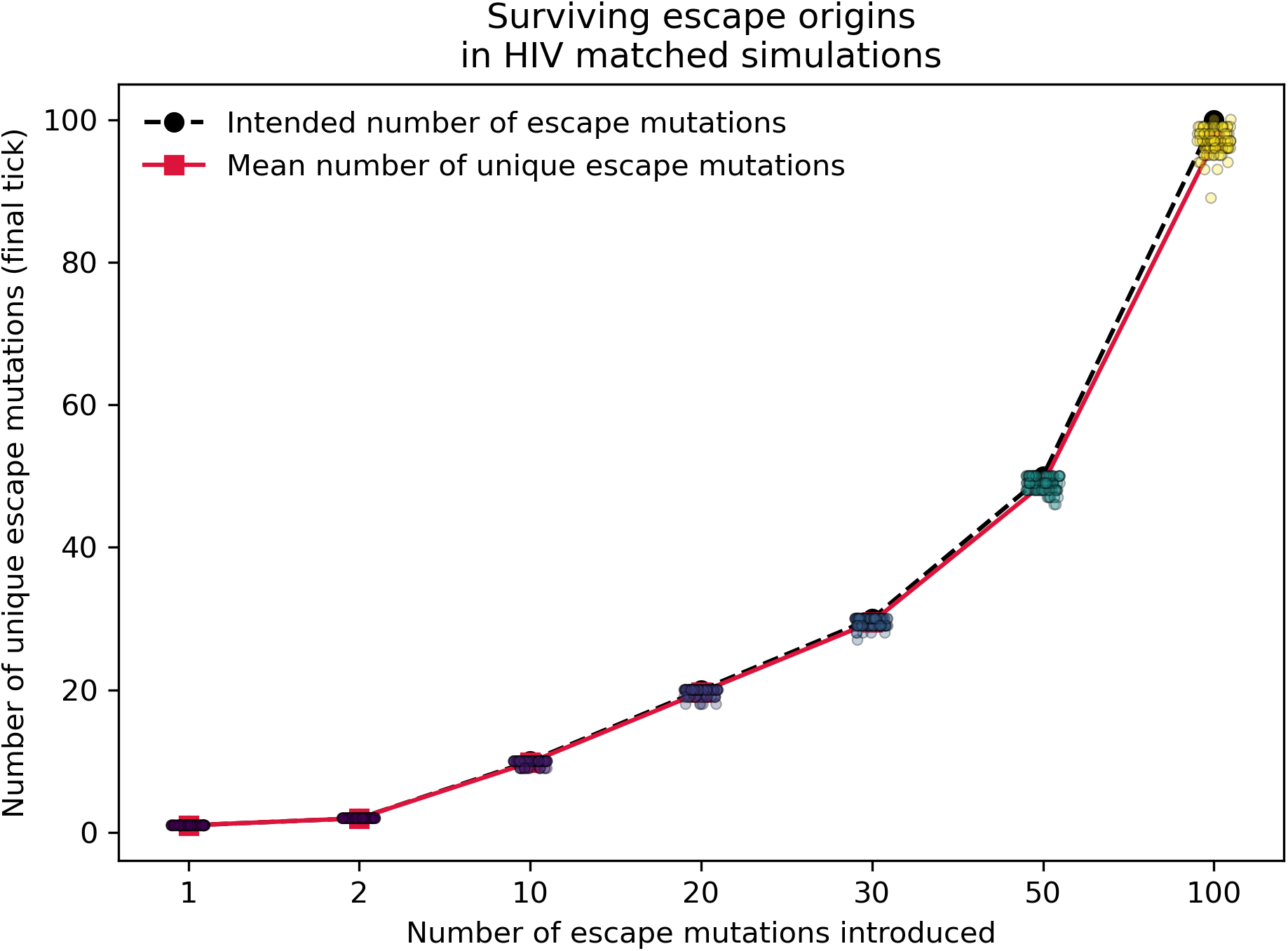
Distribution of the number of surviving origins at sweep end in HIV matched simulations. The number of surviving escape mutation origins 22 simulated generations (*≈* 8 weeks) after they were introduced. Simulations starting with 1, 2, 10, 20 30, 50, or 100 origins are included with 100 replicates shown for each scenario. Only simulations where the actual number or origins fell above a specified minimum number of origins and where the origins reached a cumulative frequency *>* 0.99 by the end of 22 generations were retained (See Materials & Methods).

**Figure S5:**
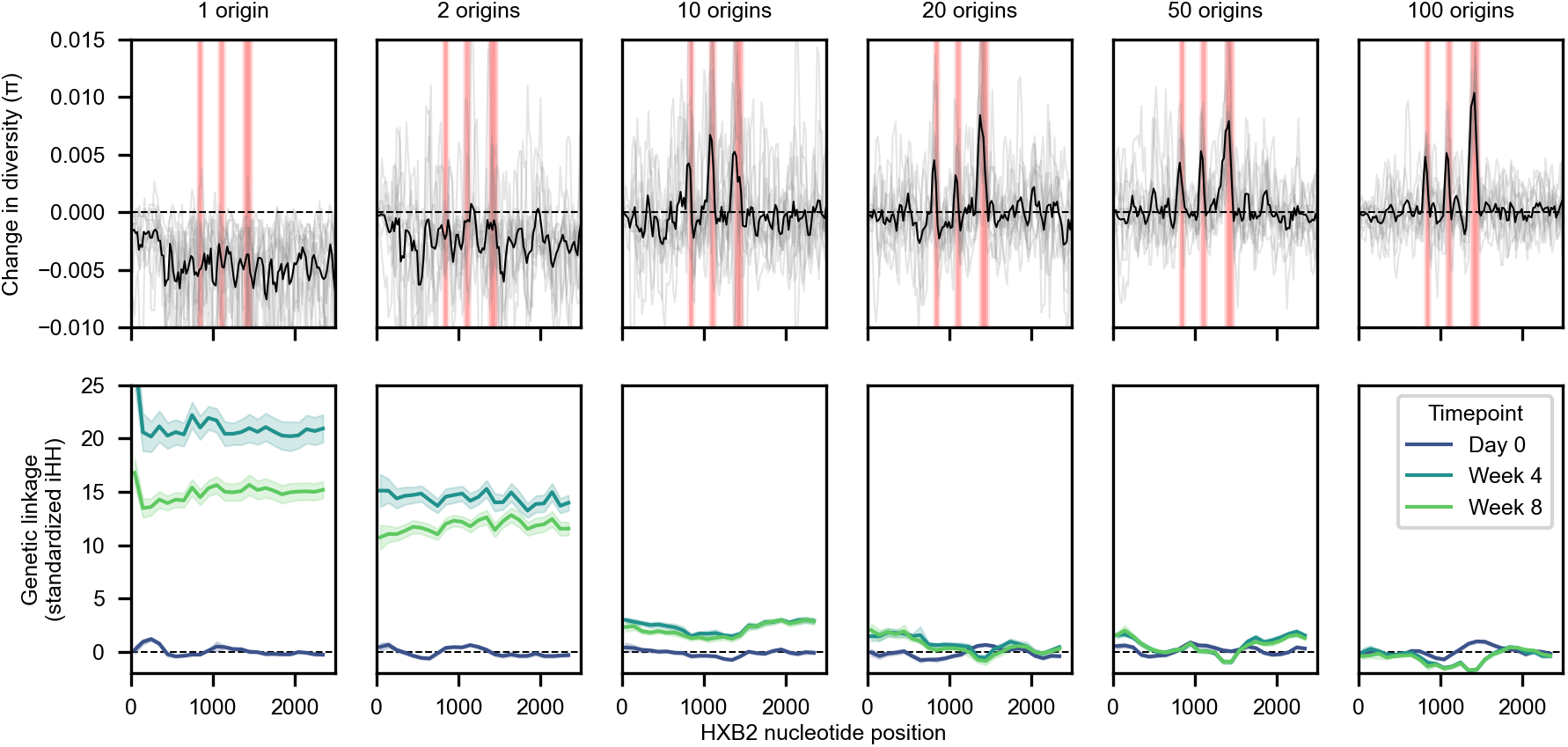
Simulated sweeps of alleles at multiple escape loci. Linkage and diversity changes are shown for sweeps of escape mutations spread across three different possible escape regions, indicated via red shading in the top panels. Sequences were limited to carrying a single beneficial escape mutation. **(Top)** Change in average pairwise Hamming distance between trial “day 0” and trial “week 4” (11 simulated generations later) for sweeps originating from 1 to 100 highly beneficial mutations introduced into randomly selected HIV genomes at a given escape locus within these three regions (see Materials & Methods). Individual simulation replicates are shown in light gray while the mean over 10 replicates is shown in black. Hamming distance is calculated in 60 nucleotide bins advanced by 20 nucleotides. The escape loci are shaded red. **(Bottom)** Standardized integrated haplotype homozygosity (iHH) plotted versus HXB2 nucleotide coordinate during simulated selective sweeps of different numbers of origins (Materials & Methods). Data are pooled across 10 replicates and plotted as a binned average (100 nucleotide positions per bin) with 95% confidence intervals on each bin.

**Figure S6:**
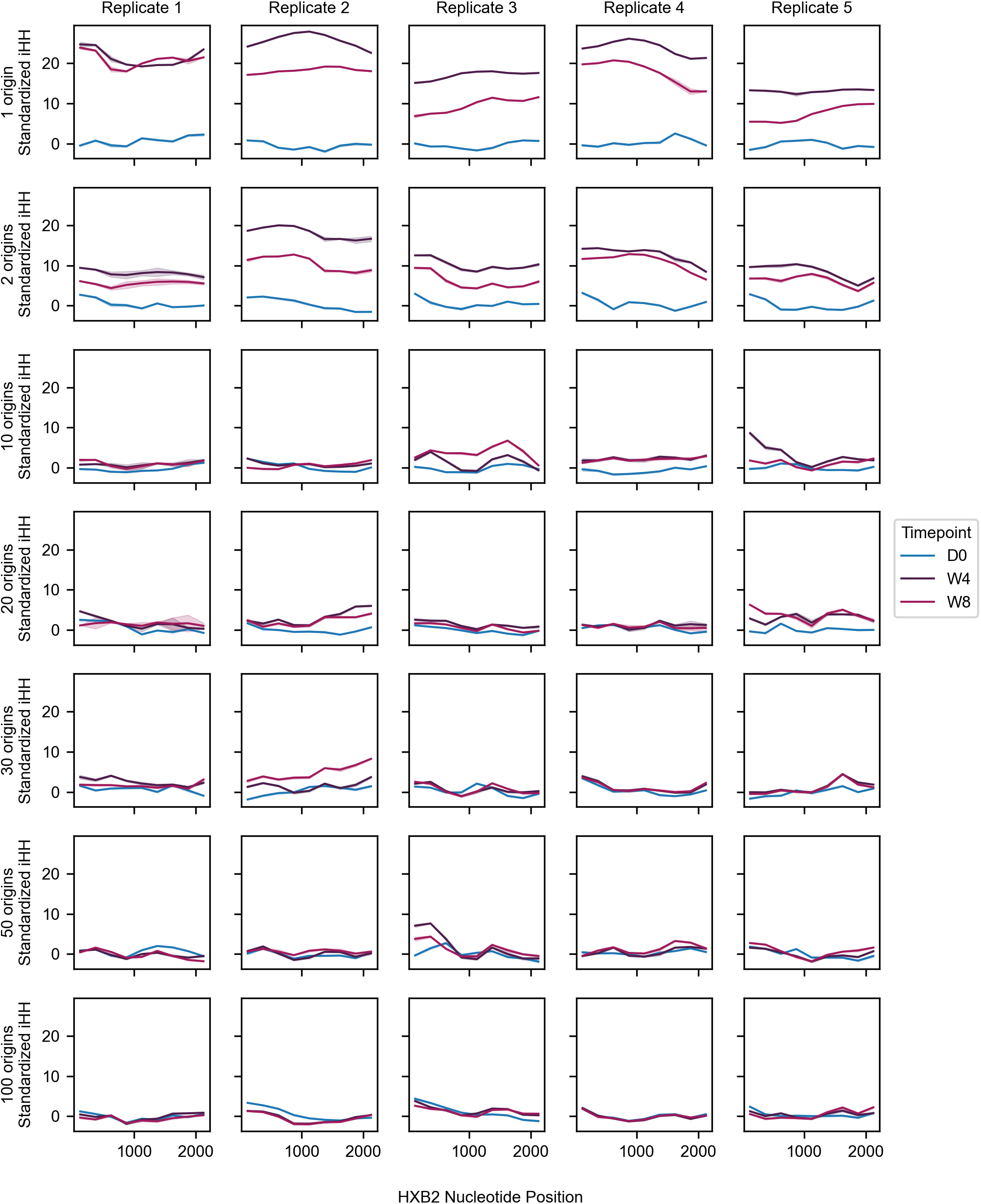
Standardized iHH over time in single simulated replicates. Standardized integrated haplotype homozygosity (iHH) is plotted versus HXB2 nucleotide coordinate during simulated selective sweeps (Materials & Methods). Each panel contains data for a single replicate of a simulation scenario with a given number of sweep origins. Five randomly selected examples are shown for each condition. Data are plotted as a binned average (100 nucleotide positions per bin) with 95% confidence intervals on each bin 3s1hown via shading.

**Figure S7:**
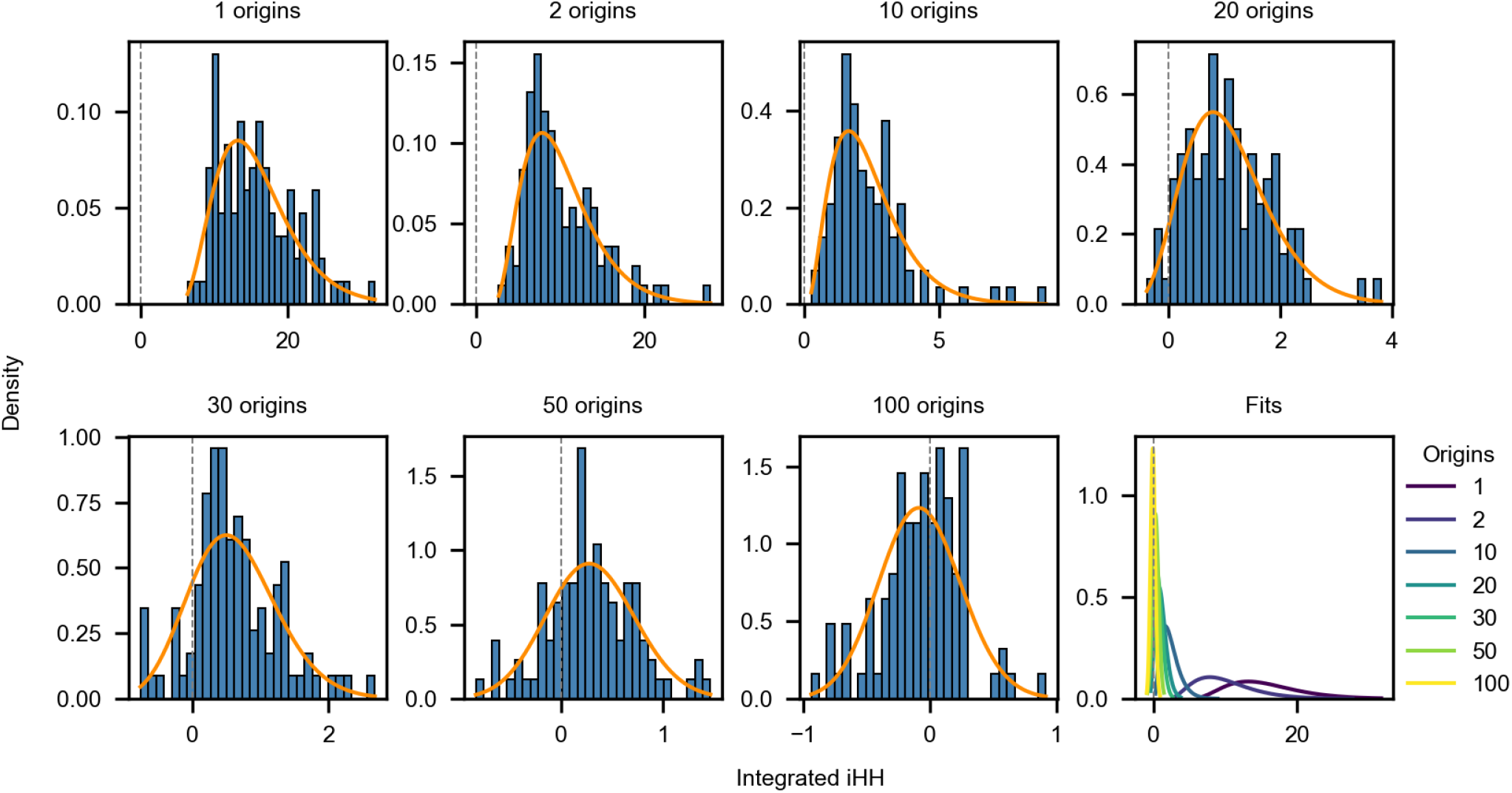
Density plots of integrated iHH for simulated sweeps with set numbers of origins. Selective sweeps simulated under HIV matched conditions are shown. Each density plot shows data from 100 replicate sweeps with a specific number of introduced origins. Orange curves indicate the gamma distribution fit to the corresponding simulated data (Table S1). The final panel shows the gamma distributions across all origin numbers for comparison.

**Figure S8:**
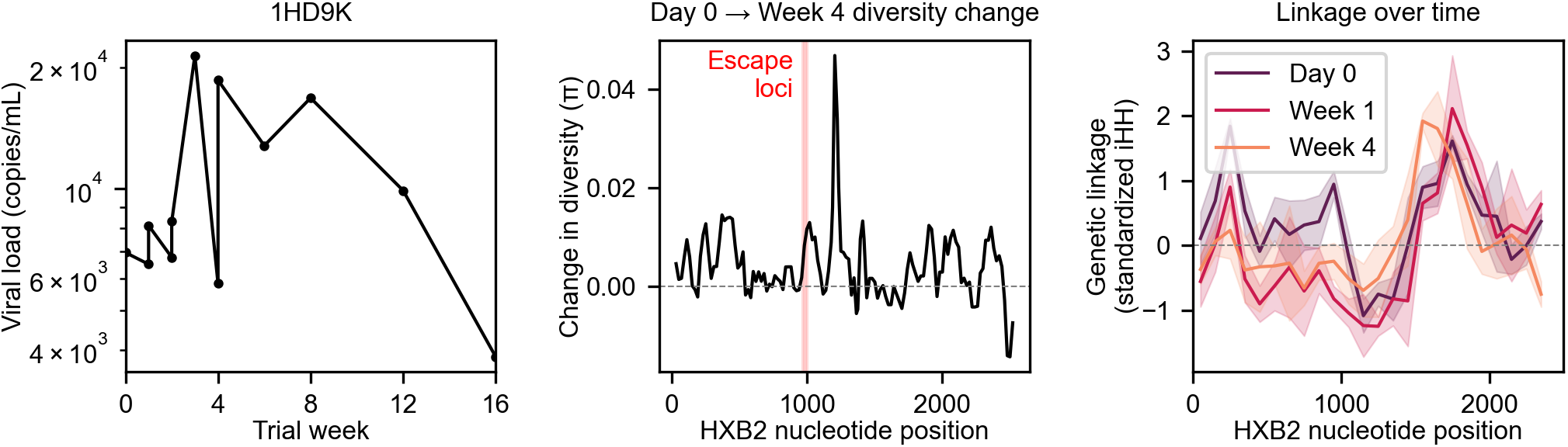
Viral load, diversity, and iHH in non-responder 1HD9K during the 10-1074 trial. **(A)** Viral load trajectory in 1HD9K throughout the 10-1074 trial. **(B)** Change in diversity in participant 1HD9K, measured via average pairwise Hamming distance between trial day 0 and trial week 4. **(C)** Standardized iHH is plotted versus HXB2 nucleotide coordinate for participant 1HD9K during the 10-1074 trial (Materials & Methods). Data are plotted as a binned average (100 nucleotide positions per bin) with 95% confidence intervals on each bin shown via shading.

**Figure S9:**
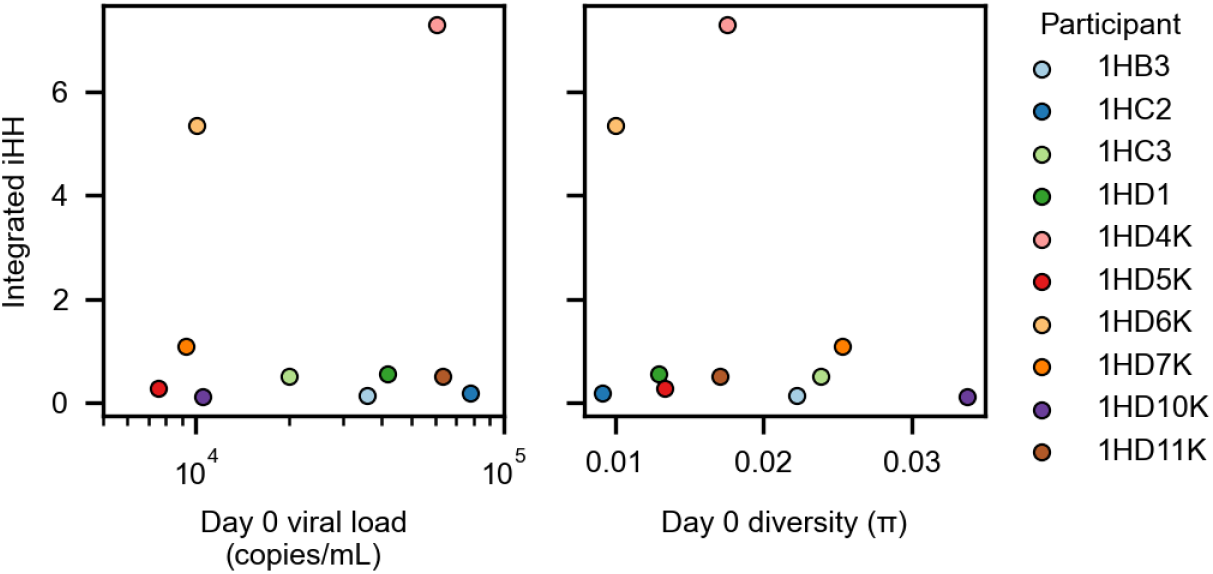
Integrated iHH (iiHH) in 10-1074 trial participants compared to pre-infusion characteristics. Integrated iHH for each trial participant is plotted in comparison to their viral load and intra-host HIV diversity pre-infusion, measured via *env* gene wide Hamming distance.

**Figure S10:**
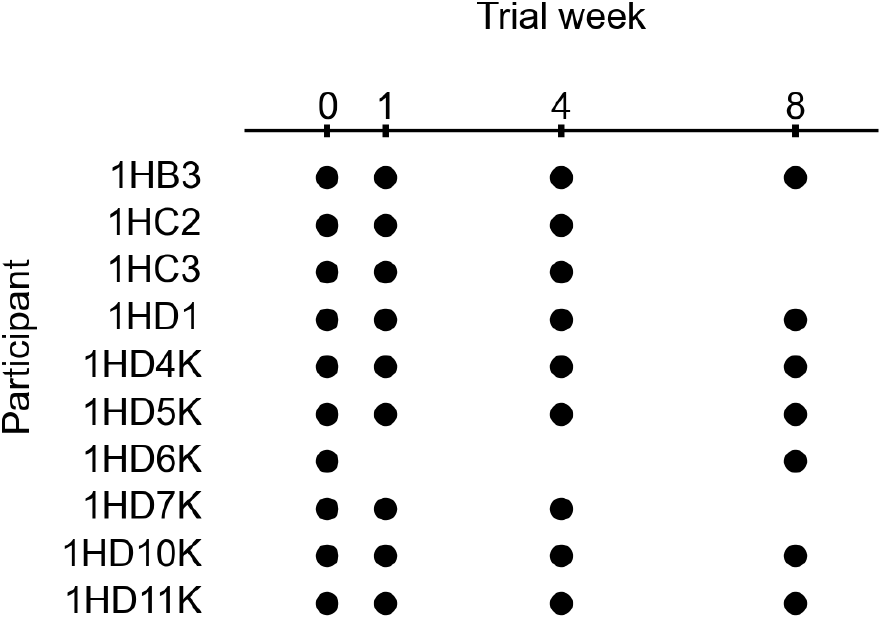
Sampling time points of 10-1074 sequences analyzed in this investigation. Dots denote all time points between trial day 0 and trial week 8 with available sequencing data.

